# MutAIverse: An AI-Powered, Mechanism-backed Platform for Discovering Novel DNA Adducts and their precursor Genotoxins

**DOI:** 10.1101/2025.08.26.672508

**Authors:** Shiva Satija, Sanjay Kumar Mohanty, Sachin Jorvekar, Anshul Verma, Saveena Solanki, Vishakha Gautam, Sakshi Arora, Aayushi Mittal, Arushi Sharma, Sonam Chauhan, Subhadeep Duari, Suvendu Kumar, Avdhesh Rai, Anupam Das, Kaberi Kakati, Kishore Das, Anupam Sarma, Juhi Tayal, Debarka Sengupta, Anurag Mehta, Roshan M Borkar, Gaurav Ahuja

**Author notes:** **Correspondence:** Gaurav Ahuja; Roshan M Borkar. Shared First Authors (Shiva Satija, Sanjay Kumar Mohanty, Sachin Jorvekar). Lead Contact: Gaurav Ahuja.

## Abstract

Genotoxin exposure leads to DNA adduct formation, potentially causing mutations if unrepaired. Current DNA adductomics platforms or analytical workflows are limited by incomplete spectral libraries, reliance on experimentally validated adducts, limited cellular contexts, and inefficient computational methodologies. We introduce MutAIverse, an advanced AI-driven DNA adductomics analysis platform that overcomes these limitations by leveraging intracellular mechanistic modeling of genotoxin bioactivation to construct a comprehensive DNA adduct library and offers advanced spectral mapping. MutAIverse integrates experimentally validated and chemically valid putative DNA adducts, enabling interpretable retrograde tracking to parental genotoxins. Validation of MutAIverse against experimental MS/MS datasets demonstrated the detection of both known and novel adducts. Furthermore, application to in-house generated DNA adductomics data from tissue biopsies of smokeless tobacco-induced head and neck cancer patients revealed selective enrichment of both novel and known DNA adducts; a subset of them was validated using MS/MS analysis. Collectively, MutAIverse provides a robust, end-to-end, and interpretable platform for advanced DNA adductomics analysis.

## INTRODUCTION

Despite its resilience as the master blueprint of cellular architecture, deoxyribonucleic acid (DNA) remains susceptible to damage from exogenous chemical genotoxins ^1^. Although cellular DNA repair mechanisms effectively recognize and repair most DNA damage, occasional failures can lead to mutations during replication or transcription ^2,3^. Electrophilic genotoxins can directly or indirectly modify nitrogenous bases by forming covalent bonds, altering their chemical properties ^4^. Other genotoxins, with neutral or weak electrophilic properties, are transformed by cellular enzymes into reactive oxygen and nitrogen species (ROS/RNS) ^5^. These highly reactive species can then bind to nitrogenous bases, disrupting the integrity and function of DNA. The vast array of chemicals in the environment, combined with the global burden of cancer, highlights the urgent need for advanced biomonitoring techniques like DNA adductomics ^6^. Importantly, DNA adductomics is a powerful and rapidly evolving technique that has successfully linked specific environmental genotoxins to particular cancers. For example, exposure to polycyclic aromatic hydrocarbons (PAHs) is associated with lung cancer, aromatic amines with bladder cancer, and aflatoxin B1 with liver cancer ^7,8^. These findings demonstrate the potential of DNA adductomics to identify carcinogens, elucidate their mechanisms of action, and improve our understanding of cancer development.

To date, various methods have been used to biomonitor DNA adducts in biospecimens, including ^32^P-post labeling, immunoassays, fluorescence labeling, and mass spectrometry-based approaches ^5^. Advancements in nanopore sequencing, especially Oxford Nanopore Technology (ONT) ^5,9^, allow for the detection and characterization of DNA adducts at single-nucleobase resolution, preserving their native forms and positional context ^9^. High-Resolution Mass Spectrometry (HRMS) has significantly advanced the field by offering superior sensitivity and accuracy in detecting and characterizing a wide range of DNA adducts, enabling researchers to identify specific chemical exposures and their biological consequences ^4,5,10^. Due to the vast diversity of carcinogens and genotoxins and the resulting complexity of DNA adducts, advanced analytical techniques are essential for their purification, identification, and characterization ^11,12^. While DNA adducts vary in stability; some are readily detected in biospecimens, while others are unstable, decomposing spontaneously at body temperature or undergoing rapid repair ^13^. For example, alkylation of purine bases often destabilizes the glycosidic bond, leading to apurinic sites. In contrast, pyrimidine adducts tend to be more stable, making them preferable biomarkers due to their abundance and ease of analysis ^5^. Mass

spectrometry (MS) has become a crucial tool for identifying DNA adducts, but its efficacy is hampered by several challenges ^14,15^. First, existing DNA adduct-specific spectral libraries are limited in scope. Widely used databases like the Human Metabolome Database (HMDB) ^16^, METLIN ^17^, MassBank ^18^, and the Kyoto Encyclopedia of Genes and Genomes (KEGG) ^19^ are not designed for DNA adductomics and contain few DNA adduct entries. While dedicated resources like dnaadductportal ^20^ and The DNA adduct database ^14^ have emerged, identifying novel adducts remains difficult. This highlights the need for a comprehensive DNA adduct spectral library to improve detection capabilities and advance our understanding of adduct structural diversity ^15^. Second, standard DNA adductome analysis lacks robust computational methods to connect both known and unknown adducts to their parental genotoxins ^4^. This limitation, coupled with the restricted scope of existing MS spectral libraries, hinders the identification of potential adducts. Finally, traditional screening methods for MS-based DNA adductomics datasets often lack accuracy and explanatory power.

To address these challenges, we introduce MutAIverse, an advanced AI-driven platform that overcomes the limitations of conventional approaches. Unlike traditional methods that rely on limited experimental data, MutAIverse utilizes intracellular simulation modeling of genotoxin metabolism and bioactivation, incorporating factors like oxidative stress and electrophilicity. This approach generated a comprehensive, end-to-end spectral mapping workflow for thorough identification of DNA adducts in biospecimens. We rigorously validated the MutAIverse spectral library and its computational workflow using diverse public and in-house DNA adductomics datasets. Taken together, MutAIverse is a standalone Python package that provides an extensive MS/MS-based DNA adduct spectral library, streamlined analytical workflows, and sophisticated computational tools for accurately identifying DNA adducts and their parental genotoxins.

## RESULTS

### *In silico* Modeling of Genotoxin Metabolism Reveals a Vast Landscape of Potential DNA Adducts

Accurate and comprehensive identification of DNA adducts from MS/MS spectra requires a robust reference library with extensive coverage. Existing libraries, limited by reliance on experimentally derived spectra, contain fewer than 400 entries. In contrast, computational approaches, especially those using generative AI, can leverage DNA adduct-associated chemical properties to predict novel DNA adducts. However, these approaches are limited by the scarcity of training data (fewer than 400 known adducts), hindering their ability to capture the full chemical diversity of DNA adducts. To overcome this limitation, we developed a novel approach independent of experimentally validated DNA adducts. We introduce MutAIverse, an AI-powered, mechanism-based platform that provides a comprehensive workflow for identifying both known and novel DNA adducts from MS/MS spectra **(Figure 1a)**. MutAIverse operates on the principle that modeling the bioactivation and biotransformation of potential adduct-forming genotoxins enables the generation of chemically valid DNA adducts. Briefly, chemical genotoxins can either directly form DNA adducts or undergo intracellular phase I and II metabolism, producing reactive or non-reactive intermediates. Reactive intermediates, such as electrophiles, can then form covalent bonds with DNA bases, resulting in adducts. This comprehensive modeling approach enables MutAIverse to accurately simulate diverse pathways of DNA adduct formation and identify a wide range of adducts. We began by compiling a list of 3,288 experimentally validated genotoxins, assuming they all contribute to DNA adduct formation, either directly or indirectly **(Figure 1b)**. Chemical composition analysis revealed that these genotoxins span diverse chemical classes, primarily benzenoids or organoheterocyclic compounds, reflecting significant chemical diversity **(Figure 1c)**. We then used Biotransformer 3.0 ^21^ to simulate the intracellular metabolism of these genotoxins, predicting the resulting metabolites or fragments via phase I and phase II enzyme rules. Biotransformation simulations showed that 92.2% (3,033) of the genotoxins were metabolized *in silico*. Further analysis revealed that Non-biotransformed (NBT) genotoxins occupied a similar chemical space as biotransformed (BT) genotoxins **(Figure 1d)**, except for enrichment in ether functional groups **(Figure 1e)**. Neither group showed a distinct subclass preference **(Figure 1c)**. To account for the possibility that metabolized genotoxin products may undergo additional rounds of biotransformation, given their accessibility to the cellular metabolism machinery, we performed iterative biotransformation simulations. In these simulations, the biotransformation cycle parameter, *k*, was varied from 0 (no biotransformation) to 3 (three rounds of biotransformation). These additional steps ensured that most, if not all, possibilities of genotoxin processing within the cell were considered. As expected, the number of metabolized products increased with higher *k* values, ranging from 25,274 (*k*=1) to 217,434 (*k*=3) **(Figure 1f)**. Further analysis revealed that the common biotransformations included hydroxylation, desaturation, reduction, oxidation, and epoxidation, among others **(Figure 1g)**.

**Figure 1.**
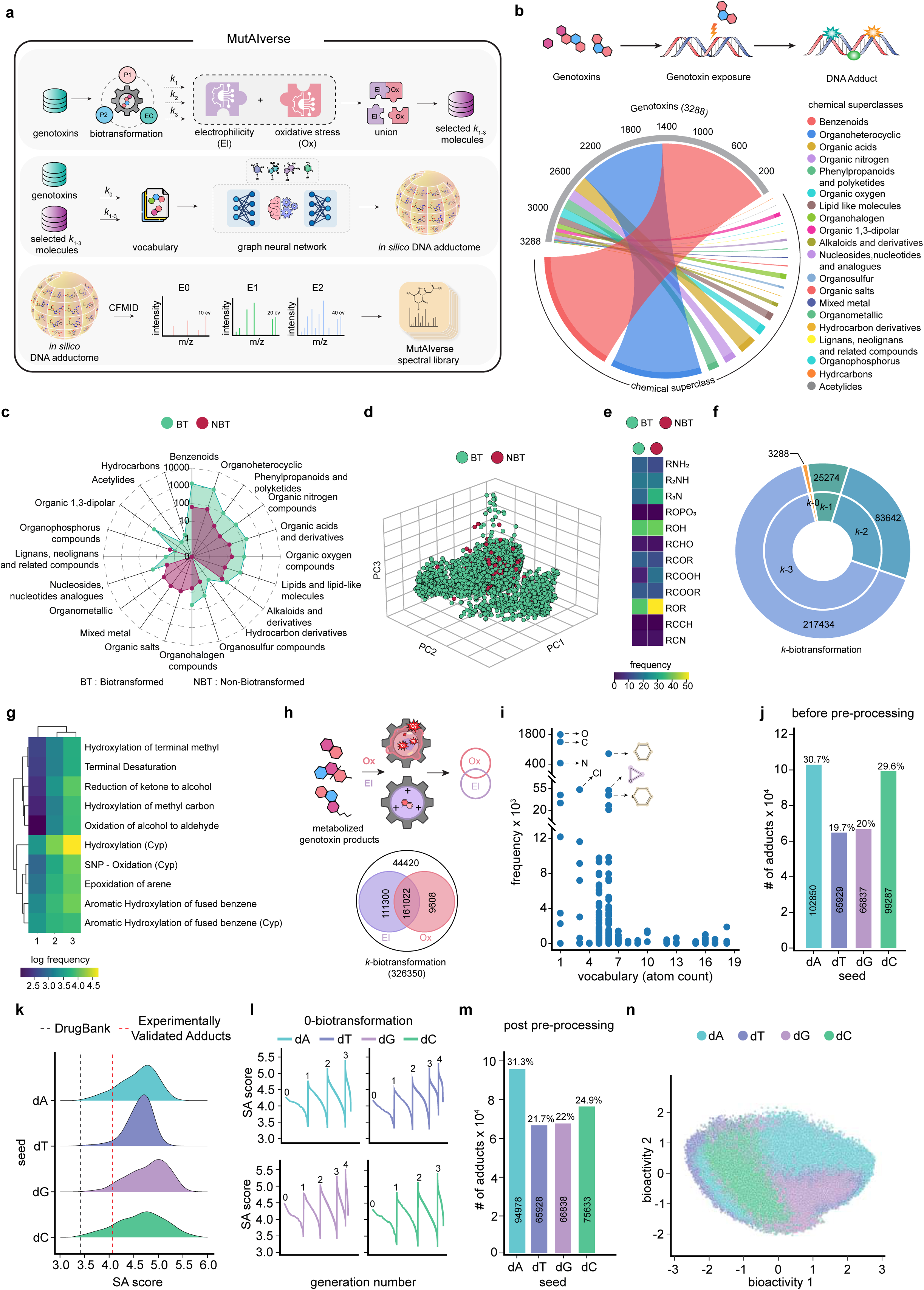

Next, to identify reactive genotoxins and their metabolites, we used two previously published machine-learning models ^22^. These models predict the potential of compounds or metabolites to induce oxidative stress or exhibit electrophilic properties. Screening the biotransformed genotoxin fragments (metabolized products) using these models revealed that 86.38% of them contribute to either oxidative stress or electrophilic properties (reactive metabolites) **(Figure 1h)**. Consequently, we selected the union of these reactive metabolites for downstream analysis under the assumption that both could equally contribute towards DNA adduct formation. Once compiled, filtered, and selected, these metabolized, potentially electrophilic, and oxidative stress-inducing genotoxin fragments were used to build a vocabulary for the downstream generation of DNA adducts. This vocabulary included abundant atoms like carbon, oxygen, chlorine, and nitrogen, as well as simple organic compounds like benzene **(Figure 1i)**. Finally, to generate a comprehensive DNA adduct library, we used the MultI-constraint MOlecule SAmpling (MIMOSA) framework ^23^. MIMOSA leverages input molecules as initial guesses and samples new molecules from the target distribution, which, in our case, are reactive genotoxin metabolized products (electrophilic and capable of inducing oxidative stress). The framework pretrains two property-agnostic graph neural networks (GNNs) for molecule topology and substructure-type prediction. Starting with DNA-specific deoxyribonucleosides: deoxyadenosine (dA), deoxycytosine (dC), deoxyguanine (dG), and deoxythymine (dT) as seed molecules, we employed the aforementioned metabolized genotoxin fragment-based vocabulary within the MIMOSA framework **(Figure 1j)**. The entire process of generating novel DNA adducts was iteratively performed using the metabolized genotoxin-generated vocabularies obtained from each biotransformation cycle parameter (*k* = 0, 1, 2, and 3). The impact of *k*-biotransformation on GNN model performance was measured by assessing the validation loss at various levels (*k* = 0-3) **(Supplementary** Figure 1a). More precisely, the validation loss was largest in the case of *k* = 0, but it significantly decreased with biotransformation (*k* ≥ 1) **(Supplementary** Figure 1a). The increase in *k* values resulted in a decrease in the variability of model performance, leading to increased learning of the GNN model, as suggested by the narrower confidence intervals **(Supplementary** Figure 1a). The generated DNA adducts were synthesized with the optimization of the Synthetic Accessibility (SA score) metric, which assesses the feasibility of chemical synthesis. The generated adducts exhibited a broad spectrum of SA scores, reflecting the varied complexity and synthesizability of the molecules **(Figure 1k, l and Supplementary Figure b-d)**. This approach yielded 303,377 unique DNA adducts, with individual contributions from each deoxyribonucleoside as follows: 94,978 from dA, 65,928 from dT, 66,838 from dG, and 75,633 from dC **(Figure 1m)**. Clustering analysis of MutAIverse adducts in bioactivity and spectral space resulted in distinct clusters showing the diversity of MutAIverse-generated DNA adducts **(Figure 1n, Supplementary** Figure 1e). Finally, to confirm the specificity of our approach, we performed similar simulations with non-genotoxic compounds (*k* = 0 to 3) and compared the resulting fragments to those generated from genotoxins using machine learning. Our results suggest that with an increasing value of *k*, the AUC-ROC values also increased with the same hyperparameters, suggesting differences in the chemical compositions of fragments obtained for genotoxins and non-genotoxins **(Supplementary** Figure 1f-i). In summary, by simulating the intracellular metabolism of genotoxins and leveraging a novel AI-driven framework, we have generated a comprehensive and chemically valid DNA adductome.

### MutAIverse Generates Novel and Known DNA Adducts While Preserving Chemical Properties

Recent efforts to compile a dedicated database of DNA adducts, accompanied by the construction of a standard reference MS/MS spectral library, have identified 279 experimentally validated (referred to as EVA, Experimentally Validated Adducts) and 303 chemically plausible DNA adducts (referred to as SA, Suspected Adducts) ^14^ **(Figure 2a)**. Next, we investigated whether the DNA adductome generated using the MutAIverse framework includes previously characterized EVA and SA DNA adducts. We first compared physicochemical properties (atom count, molecular weight, logP, TPSA, and Kier-Hall Index ^24^) between MutAIverse, EVA, and SA adducts **(Figure 2b)**. Overlapping distributions across these properties demonstrate that our mechanism-based approach captures the chemical space of known DNA adducts. We also compared topological features (kappa 1, kappa 2, kappa 3, and LabuteASA) between MutAIverse, EVA, and SA adducts. Similar results for these topological features further validate that our framework preserves key chemical and structural features of DNA adducts **(Figure 2c, Supplementary** Figure 2b). Furthermore, TMAP ^25^ clustering of MutAIverse adducts revealed distinct clusters based on properties like logP, molecular weight, and ring count **(Supplementary** Figure 2a). This analysis highlighted variations in hydrophobicity, size, and complexity, providing insights into the distribution and relationships among the adducts. To evaluate the level of structural diversity, we also performed Tanimoto similarity (TS) and Overlap coefficient analyses between the fingerprints computed for the MutAIverse adducts and those in the EVA and SA datasets. The maximum TS scores ranged from 0.4 to 1, indicating a considerable degree of structural similarity between the libraries **(Figure 2d, e; Supplementary** Figure 2f**, g)**. In contrast, the minimum TS scores ranged from 0.0 to 0.3, reflecting a broad range of structural diversity among the chemically valid DNA adducts generated by MutAIverse. These analyses demonstrate that our framework captures a wide range of structural variations within the DNA adductome. Next, we conducted a comparative analysis of the DNA adducts from MutAIverse, EVA, and SA datasets at the spectral peak level. This involved generating *in silico* ESI (+) MS/MS spectral peaks for the DNA adducts precursor [M+H]+ using the CFM-ID ^26,27^ computational framework at three distinct collision energy settings, denoted as 10eV (E0), 20 eV (E1), and 40eV (E2), corresponding to different stages of energy application and the resulting fragmentation patterns. Subsequently, we employed the *matchms* ^28^ Python library to perform pairwise comparisons of these spectral peaks, utilizing the Cosine Similarity metric to quantify the degree of correspondence between MutAIverse and EVA or SA DNA adducts **(Figure 2f-h)**. In the comparison between MutAIverse-generated adducts and EVA, we observed a pronounced peak in the Cosine Similarity distribution, with the mean (95% confidence interval) ranging from 0.94 (0.93, 0.95) **(Figure 2g)**. Detailed analysis indicated that the highest Cosine Similarity values were recorded at collision energy level E0, followed by E1 and E2 **(Figure 2g; Supplementary** Figure 2c). A comparable trend was observed in the analysis of MutAIverse-generated adducts versus SA adducts **(Figure 2h; Supplementary** Figure 2d), confirming the robustness of our approach across different datasets and energy levels. Finally, we compared the distribution of spectral features of the DNA adducts present in MutAIverse, EVA, and SA at these three collision energy levels and observed comparable spectral entropies between the groups **(Figure 2i)**. Entropy similarity analysis revealed high entropy similarity for EVA and SA adducts compared to the MutAIverse library **(Supplementary** Figure 2h). Notably, in these analyses, several MutAIverse adducts with cosine similarity = 1 were detected during the comparative analysis with EVA **(Figure 2j)**. In summary, a thorough analysis of MutAIverse-generated DNA adducts, compared with EVA and SA, demonstrated congruent distributions across the chemical, topological, and entropy metrics. This consistency underscores the fidelity of MutAIverse in replicating the chemical space and structural characteristics of known DNA adducts.

**Figure 2.**
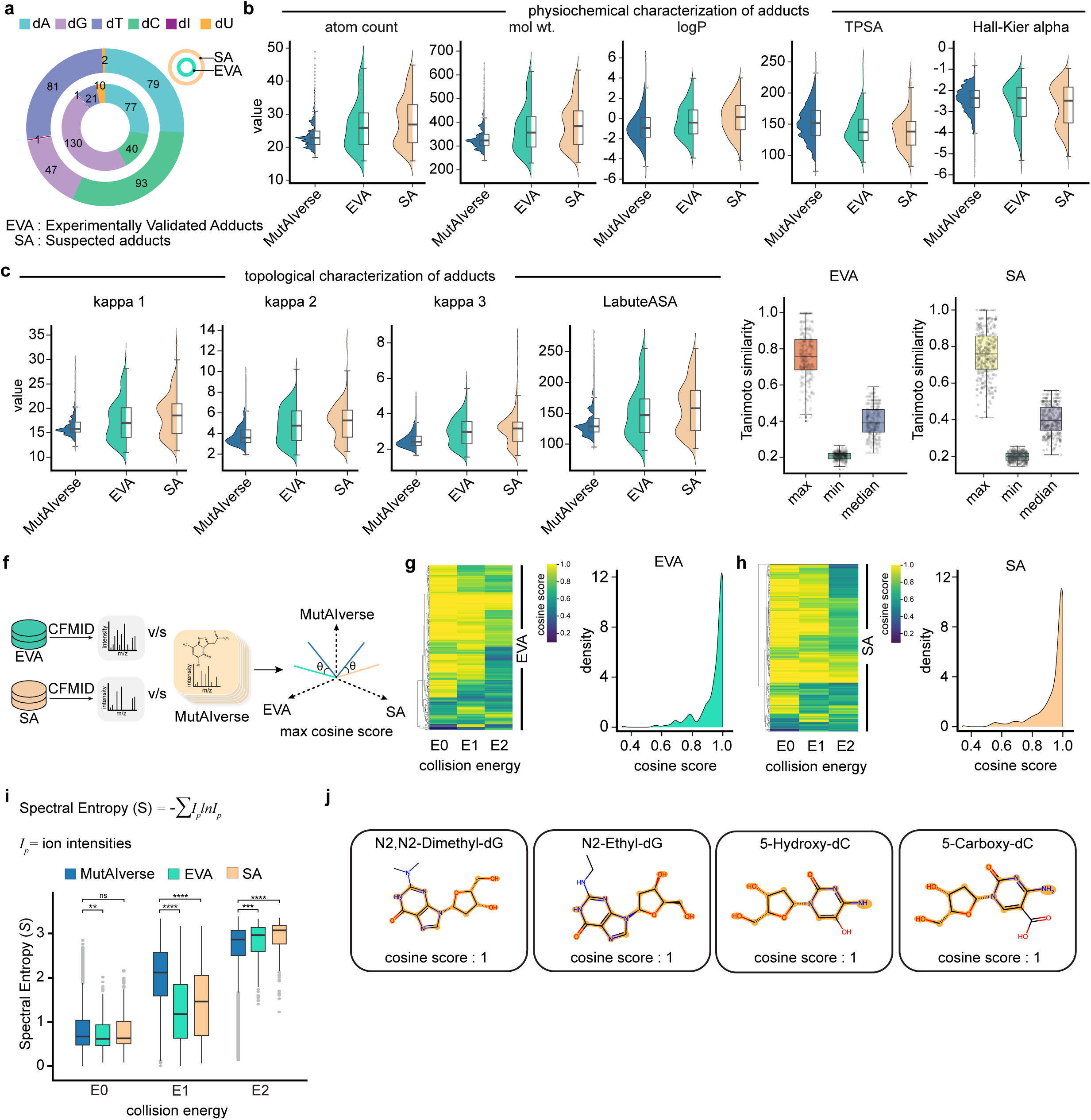

### MutAIverse framework enables Precise DNA Adduct-Genotoxin Linking

While various analytical methods excel at identifying novel DNA adducts in biospecimens, linking these adducts to their parent genotoxins remains a significant challenge. To address this, we employed an *in silico* approach that models the reversal of genotoxin bioactivation and biotransformation within cells, thereby developing methods for retrograde tracing DNA adducts back to their potential parental genotoxins. We named this sub-module as AdductLinker due to its functional importance. AdductLinker employs a multi-step computational framework: it begins by isolating the nitrogenous base from the DNA adduct to extract the genotoxin-contributing fragment via a process called splicing **(Figure 3a, b)**. It then calculates the structural fingerprints of this fragment and all genotoxin intermediates and uses *k*-nearest neighbor (kNN) clustering to identify the nearest neighboring intermediate and their corresponding genotoxins **(Figure 3a)**. Finally, AdductLinker determines the maximum common substructure (MCS) among these nearest neighbors, which is used to trace back through the genotoxin-intermediates-metabolized fragment tree, thereby identifying potential parental genotoxins **(Figure 3a; Supplementary** Figure 3a). AdductLinker outputs the queried DNA adduct, its upstream metabolized products, and the parental genotoxin, along with a confidence score. While this approach significantly accelerates the process of linking DNA adducts to their parent genotoxins, it is limited by its reliance on the genotoxins used to construct MutAIverse and the prerequisite that not all DNA adducts underwent splicing **(Figure 3c-e; Supplementary** Figure 3b). We validated and tested AdductLinker using *bona fide* DNA adduct-genotoxin pairs, including AFB-dG-Aflatoxin, N2-DB[a,h]ADE-dG-Dibenzo[a,h]anthracene, N2-DB[a,l]PDE-dG-Dibenzo[a,l]pyrene, N6-(5-Me-CDE)-dG-5-methyl-chrysene, 2-AF-dG-2-Aminofluorene and Benzo[a]pyrene-dG-Benzo[a]pyrene **(Figure 3f; Supplementary** Figure 3f). Our results suggest that the AdductLinker sub-module of MutAIverse successfully backtracked both the upstream metabolized products and the parental genotoxins, demonstrating its efficacy in identifying potential genotoxin origins for a given DNA adduct.

**Figure 3.**
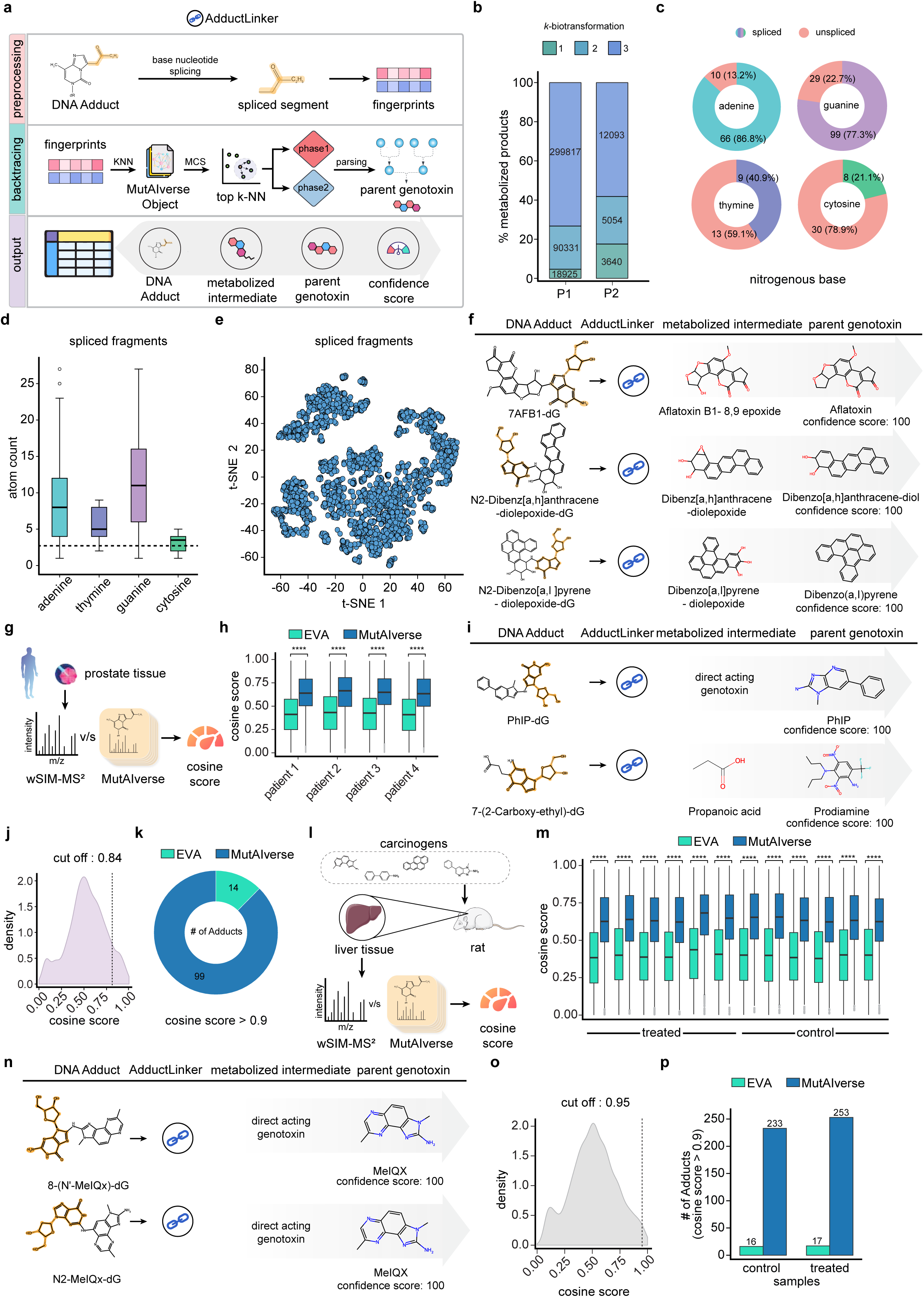

To test this sub-module further in analyzing experimental MS/MS data from biospecimens, we utilized the recently published datasets from Walmsley and colleagues ^29^. The dataset contained tumor-adjacent prostate tissue from 4 prostate cancer patients. In the original study, the authors identified 361 putative and six *bona fide* DNA adducts, including dG-C8 PhIP, N2-Dimethyldioxane-dG, 4-OH-2-heptenal-dG, (E)-4-hydroxy-2-nonenal-dC, (E)-4-oxonon-2-enal-dC and M1AA-dA using a standard DNA adduct-specific MS library which contained 349 adducts of 279 unique masses ^29^. We analyzed this dataset with the MutAIverse computational framework while selecting the matchms-based cosine similarity calculations **(Figure 3g)**. Our results identified multiple known and novel DNA adducts, including the ones reported in the original study. The six of the *bona fide* adducts identified were Phenylimidazo[4,5-b]pyridine-dG, N2-Dimethyldioxane-dG, E-4-Oxonon-2-enal-dC, (E)-4-Hydroxy-2-nonenal-dC, 3-(2′-Deoxy-β-D-ribofuranosyl)-7-methyl-8-formyl[2,1-i]pyrimidopurine-dA, and 4-Hydroxy-2-heptenal-dG. Notably, Phenylimidazo[4,5-b]pyridine-dG was a reported DNA adduct and was mapped to an experimentally validated adduct (EVA) of the MutAIverse with a high cosine similarity of 0.84. Furthermore, the LinkerAdduct sub-module mapped this adduct to parental genotoxin phenylimidazo[4,5-b]pyridine with confidence scores of 100% **(Figure 3i)**. Similarly, in the case of N2-Dimethyldioxane-dG, another EVA, which did not match with any known spectra in the MutAIverse library but was backtracked to multiple genotoxins such as (1R,6S)-7-oxabicyclo[4.1.0]heptane and 2-methyl-2-(oxiran-2-yl)oxirane with confidence scores of 87.5%. Similar analysis with other *bonafide* adducts led to their identification in the MutAIverse library with varied cosine similarities, as well as parental genotoxins. As stated above, we also observed multiple novel DNA adducts exhibiting a CS higher than that of PhIP (0.84) **(Figure 3h-k; Supplementary** Figure 3g-j). Implementation of AdductLinker with these novel DNA adducts (CS > 0.9) suggested (2R,6R)-2,6-dimethylmorpholine, 2-[2-(1H-indol-3-yl)ethylamino]oxane-3,4,5-triol and 2-methylidenehexanal as top three potential parental genotoxins.

Next, we also used an independent, controlled rodent-based dataset. In this dataset, rats were exposed to a cocktail of genotoxic mutagens (B[a]P, 4-ABP, PhIP, AαC, and MeIQx), and post-exposure, DNA adductomics was performed on genomic DNA from liver tissue using wSIM-MS/MS mass spectrometry **(Figure 3l)**. Next, we mapped the experimentally elucidated MS/MS peaks onto a standard DNA adduct library containing EVA and compared it to the subset of the MutAIverse library containing only *in silico-*generated adducts **(Figure 3m; Supplementary** Figure 3k-m). Comparison to EVA library reproduced six *bonafide* DNA adducts: phenylimidazo[4,5-b]pyridine-dG, dG-C8-2-Amino-9H-pyrido[2,3-b]indole, dG-C8-3,8-dimethylimidazo[4,5-f]quinoxalin-2-amine, dG-C8-4-aminobiphenyl, N-(2′-deoxyadenosin-8yl)-4-aminobiphenyl, N-(2′deoxyguanosin-N2-yl)-4-aminobiphenyl as mentioned in the original study.

Notably, Phenylimidazo[4,5-b]pyridine-dG, an EVA, demonstrated a cosine similarity of 0.76 and was backtracked to phenylimidazo[4,5-b]pyridine by AdductLinker sub-module as a potential parental genotoxin with confidence scores of 100 **(Figure 3i)**. The adduct dG-C8-2-Amino-9H-pyrido[2,3-b]indole showed a cosine similarity of 0.90 and was matched to the genotoxin 2-Amino-9H-pyrido[2,3-b]indole (AαC) with confidence scores of 100 **(Figure 3n)**. A similar analysis with other *bonafide* adducts resulted in their identification within the MutAIverse library, exhibiting varied cosine similarities and corresponding parental genotoxins. Similar to the prostate cancer dataset, we also observed multiple novel DNA adducts when mapping with the MutAIverse library **(Figure 3o, p)**. Taken together, all these results collectively underscore the robustness, comprehensiveness, and accuracy of the MutAIverse workflow in effectively identifying and validating known and novel DNA adducts within complex MS/MS datasets.

### MutAIverse Identified Novel DNA Adducts in Smokeless Tobacco-Exposed Head and Neck Cancer

Identifying novel DNA adducts with MutAIverse advances our understanding of adduct formation and holds promise for clinical translation. This could facilitate the development of new diagnostic biomarkers and therapies. As proof of concept, we analyzed head and neck tumor biopsies and adjacent non-malignant tissues from patients with a history of smokeless tobacco use **(Figure 4a)**. The study included a balanced representation of both sexes **(Figure 4b)**. We performed high-resolution MS-based DNA adductomics on genomic DNA from these samples, comparing results between tumor and adjacent non-malignant tissues **(Figure 4a)**. We identified a greater number of DNA adducts with higher cosine similarity (CS) using the MutAIverse library compared to the EVA library **(Figure 4c)**. This suggests that MutAIverse covers a broader chemical space of DNA adducts. Even with a stringent CS threshold (> 0.9), MutAIverse identified significantly more potential adducts **(Figure 4c; Supplementary** Figure 4a,b). This trend was consistent across both tumor and adjacent normal tissues **(Figure 4c)**. Given the patients’ clinical histories of smokeless tobacco use, we employed the AdductLinker module of MutAIverse to unveil the potential parent genotoxins or carcinogens. By projecting the tumor-specific DNA adducts identified through MutAIverse and performing retrograde tracking with the AdductLinker sub-module, we pinpointed several notable adduct-genotoxin pairs, such as 8-(N’-4-Chloroaniline)-dA-4-chlorobenzene-1,2-diamine and N6-Dibenz[a,h]anthracene-diolepoxide-dA, and 3,4-dihydronaphtho[1,2-b]phenanthrene-3,4-diol. Other alkylated DNA adducts such as 3-methyl-dA, N6-methyl-dA, and 3-ethyl-dA, along with ethno adducts 3,N4-etheno-dC were also detected. These findings suggest that the genotoxic agents identified are likely implicated in malignancy, as the adducts 3-methyl-dA and N6-Methyl-dA were specific to tumor tissues and detected in 18.51% of the samples **(Figure 4d)**. The spectral search of EVA against the MutAIverse library revealed a substantial degree of similarity **(Figure 4e)**, with an average cosine similarity of 0.94. MutAIverse DNA adducts were detected across both tumor and tumor-adjacent tissues (referred to as normal), while 31 MutAIverse DNA adducts were exclusive to tumor tissue **(Figure 4f, g)**. Chemical similarity analysis revealed the structural divergence between EVA and adducts created by MutAIverse **(Figure 4h, i)**, indicating that MutAIverse covers a wide range of chemical variations between EVA and MutAIverse adducts detected in tumor and tumor-adjacent samples. The AdductLinker predicted Ketene as the parental genotoxin of the MutAIverse adduct with m/z 266.0739 with a confidence score of 100%. Backtracing with AduuctLinker, the MutAIverse adduct with m/z 286.1033 to hydroperoxyethane, and the MutAIverse adduct (m/z 546.2135) with Benzo[c]phenanthrene 3,4-dihydrodiol-1,2 epoxide (B(c)Phde), a well-known tobacco-related carcinogen ^30^ **(Figure 4j)**. The intensity of both validated adducts with m/z 266.0739 (MutAIverse adduct) and m/z 377.1868 (N4-Pyridyl-oxobutyl-dC) was detected in both tumor and tumor-adjacent cells **(Supplementary** Figure 4c,d). Correlation analysis of the intensity of m/z 266.07 with the age of patients across tumor and tumor-adjacent normal cells showed a weak positive correlation, whereas the intensity of m/z 377.1868 showed a negative correlation in the case of tumor-adjacent normal cells **(Supplementary** Figure 4c). A previous study involved urinary DNA adductomics analysis using high-resolution mass spectrometry on smokeless tobacco users, non-users, and head and neck cancer patients ^31^. The method was validated with synthetic DNA adduct standards, including NDAMPP, 6O-MG, 8-NG, 7-MG, 3-MG, and 8-OHdG, by matching retention times in the samples. Additional DNA adducts were identified using a database library. Statistically significant DNA adducts were confirmed using LC-ESI-QTOF-MS/MS fragmentation for detailed structural analysis. To eliminate the possibility of detecting deoxyribonucleases in the samples, an *in silico* MS/MS analysis of deoxyadenosine (dA), deoxythymidine (dT), deoxyguanosine (dG), and deoxycytidine (dC) was performed and compared with MutAIverse library. The comparative analysis revealed that MutAIverse adducts exhibited higher cosine similarity than the dA, dT, dC, and dG, further supporting the specificity of the identified adducts **(Supplementary** Figure 4f).

**Figure 4.**
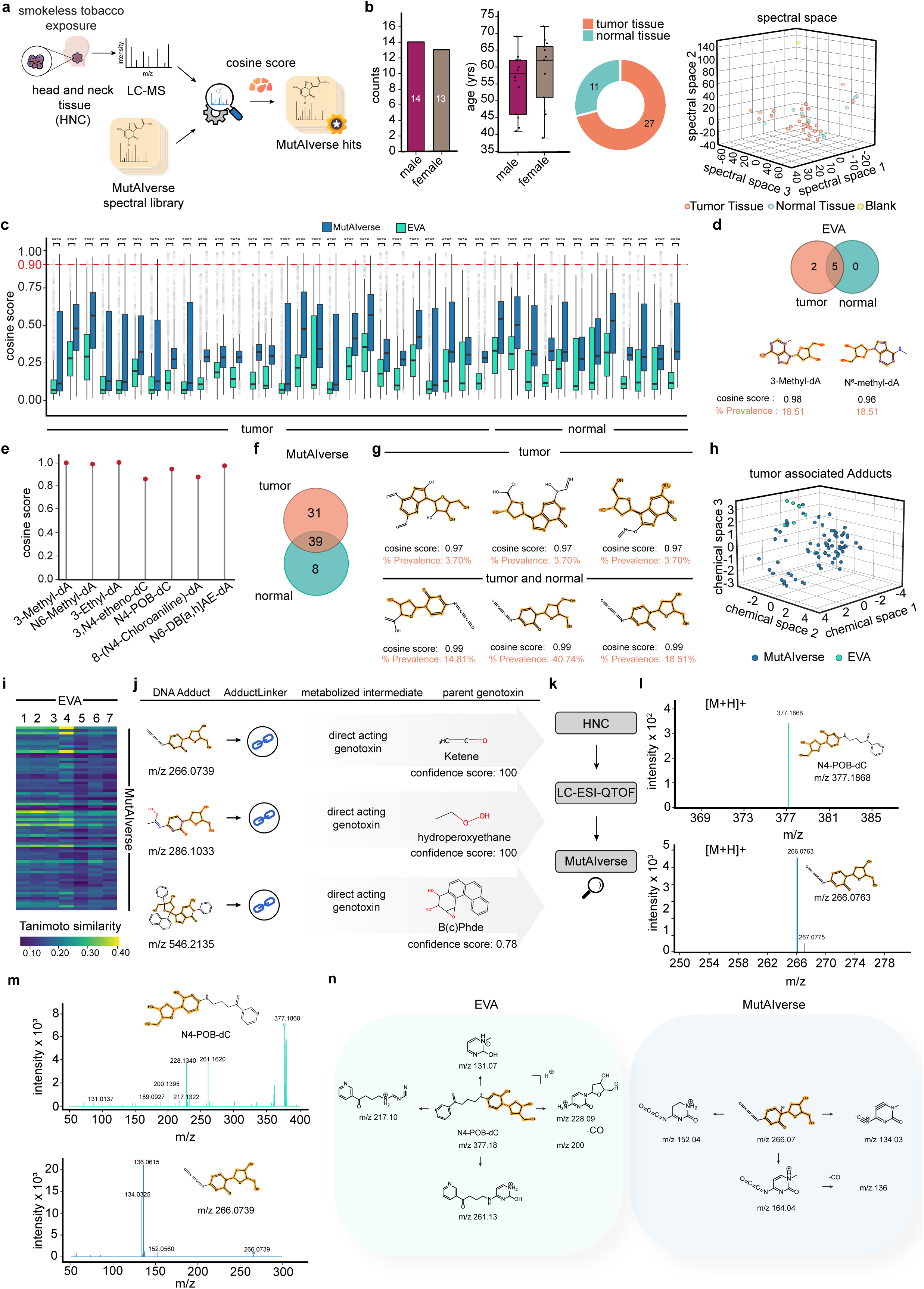

Although MutAIverse offers a comprehensive collection of genotoxin-guided and chemically valid DNA adducts, some of these adducts, while computationally and conceptually plausible, may not always be present in real-world biological scenarios. To validate the existence of novel DNA adducts identified by MutAIverse in these samples, we conducted rigorous, high-resolution MS analysis on two DNA adducts, designated N4-Pyridyl-oxobutyl-dC (N4-POB-dC) with m/z 377.1824 which is formed by the exposure of tobacco-specific nitrosamines ^10^ and a novel DNA adduct with m/z 266.0739 and MS/MS analysis of these compounds were done using low-resolution mass spectrometry **(Figure 4k-m)**. Some of the predicted fragments have mass error > 10 ppm due to their low abundance **(Figure 4n)**. Detailed examination of the MS/MS spectra confirmed the presence of the novel DNA adduct in 14.81% of tumor samples and 27.27% of tumor-adjacent control tissues. This validation not only demonstrates the efficacy of MutAIverse in uncovering novel DNA adducts but also underscores its potential as a tool for biomonitoring clinically relevant, adducts-based biomarkers associated with smokeless tobacco-related head and neck cancers.

## DISCUSSION

The identification and characterization of DNA adducts are crucial for understanding their roles in carcinogenesis and mutagenesis. Traditional methods, reliant on experimentally validated data, are limited to cataloging fewer than 400 DNA adducts, which restricts the exploration of their full chemical diversity. Computational approaches, particularly those utilizing generative AI frameworks, offer promising solutions for broadening the scope of DNA adduct identification. In this study, we introduce MutAIverse, a novel mechanism-driven generative AI framework designed to (a) generate all possible chemically valid DNA adducts from experimentally validated genotoxins, (b) provide a comprehensive MS spectral library and associated toolkits for mapping DNA adductomics datasets, and (c) support the computational prediction of parental genotoxins from detected DNA adducts. Unlike traditional methods that rely on limited experimentally validated adducts, MutAIverse employs a novel simulation-based approach to generate a more extensive array of DNA adducts.

The chemical diversity of DNA adducts generated by MutAIverse is contingent upon the selection of genotoxins used in the initial setup. Although our study incorporates an updated list of experimentally validated genotoxins, the framework is adaptable and can accommodate newly identified genotoxins as they become available. Incorporating novel genotoxins could further expand the DNA adduct library and enhance the linkage of newly generated DNA adducts to their respective parental genotoxins. Additionally, the selection of biotransformation prediction tools plays a critical role in determining the chemical diversity of the DNA adducts generated by MutAIverse. We employed Biotransformer 3.0 ^21^, a sophisticated tool that integrates both phase I and phase II metabolism predictions with EC-based metabolism and machine learning techniques. This choice contrasts with other tools such as Sygma ^32^, which relies on predefined reaction data and rule-based methods, or Gloryx ^33^, which combines rule-based approaches with machine learning. Given that genotoxins can contribute to adduct formation either directly or following biotransformation, our approach included multiple iterations of biotransformation simulations to capture a comprehensive range of possible adducts. To prioritize metabolized genotoxin fragments based on their potential for adduct formation, we utilized predictive models for electrophilicity and oxidative stress. However, additional factors could be considered to refine the adduct library further, including other reactivity types, lipophilicity, intracellular stability of genotoxin or its metabolized fragments, bioavailability, and the stability of the formed adducts. Integrating these parameters could enhance the accuracy and relevance of the DNA adductome generated by MutAIverse.

MutAIverse, while offering significant advancements in DNA adduct identification, also has several limitations that must be acknowledged. Firstly, the framework’s reliance on an initially selected set of genotoxins may result in an incomplete representation of both the number and chemical diversity of generated DNA adducts. Specifically, many chemical compounds that are not genotoxic in their native state can become genotoxins through bioactivation by cellular enzymes. Currently, MutAIverse does not model this process, which limits its ability to predict adducts formed from such transformed compounds. Integrating a genotoxicity prediction model upstream could address this issue, but it may also impact the subsequent performance and usability of the AdductLinker sub-module. Secondly, the biotransformation predictions incorporated in MutAIverse are predominantly based on human-specific metabolic pathways. This approach does not account for biotransformations that occur due to the influence of the microbiome, which can significantly affect the formation and diversity of DNA adducts, as per recent reports ^34,35^. Incorporating microbiome-based biotransformation predictions could substantially broaden the scope and chemical diversity of the generated DNA adduct library. Thirdly, the current computational framework for generating DNA adducts has been optimized using the Synthetic Accessibility (SA score) score alone. While this parameter is useful, the addition of other metrics, such as LogP (partition coefficient) and other relevant chemical properties, could further enhance the accuracy and breadth of the DNA adduct library. Parallel optimization using these additional parameters could yield a more comprehensive representation of potential DNA adducts. Fourthly, we utilized CFM-ID ^26,27^ for the *in silico* generation of ESI MS/MS spectral libraries. As a predictive tool, CFM-ID may introduce certain biases, which could affect the accuracy of the mapping algorithms and the quality of the downstream results. Lastly, although MutAIverse facilitates the identification of novel DNA adducts, their functional relevance remains to be determined. It is essential to assess whether these adducts act as transcription or replication blockers or how they contribute to mutagenesis. These levels of predictions are not presently offered by MutAIverse. All these potential biases underscore the need for further refinement and validation of the predictive models used in MutAIverse. Addressing these limitations by integrating additional models and optimizing parameters could enhance the capabilities and applications of MutAIverse, providing a more robust tool for DNA adductomics research.

Despite these limitations, MutAIverse presents several notable advancements and stands as the first proof of concept for employing a generative AI and metabolism simulation-based strategy to generate a comprehensive array of potential DNA adducts from selected genotoxins. The framework’s innovative approach marks a significant departure from traditional methods and demonstrates the potential of integrating AI with biochemical simulations. It is strongly recommended to use MutAIverse in conjunction with established MS spectral libraries, such as those provided by the National Institute of Standards and Technology (NIST) ^36^, The Human Metabolome Database (HMDB) ^16^, METLIN ^17^, MassBank ^18^, or the Kyoto Encyclopedia of Genes and Genomes (KEGG) ^19^. Additionally, adduct-specific libraries, including dnaadductportal ^20^ and The DNA Adduct Database ^14^, should be considered. Integrating these spectral libraries with MutAIverse could greatly enhance the accuracy and comprehensiveness of DNA adduct identification.

## METHODS

### Datasets Compilation

A comprehensive dataset of genotoxins, consisting of 3,288 compounds, was assembled from multiple sources, including peer-reviewed literature and established databases such as Tox21^37^. For each genotoxin, Simplified Molecular-Input Line-Entry System (SMILES) notations were retrieved through the PubChem Identifier Exchange Service. To ensure consistency and accuracy in molecular representation, the retrieved SMILES notations were standardized to canonical SMILES using OpenBabel version 3.1.1 ^38^. This standardization process is crucial for maintaining uniformity in subsequent computational analyses. For the DNA adducts, data were sourced from the DNA adductomics database ^14^, which includes a curated collection of 279 experimentally validated DNA adducts (EVA) and 303 suspected DNA adducts (SA). Similar standardization procedures for chemical representation were applied to these adducts to ensure consistency and comparability in the analysis.

### Chemical taxonomy classification

ClassyFire ^39^ was used to analyze the chemical diversity of genotoxin molecules through a structured classification approach. Initially, SMILES notations for the genotoxins were converted into Structure Data File (SDF) format using OpenBabel to ensure uniformity in data representation. The SDF files were then processed by ClassyFire, a web server that classifies chemical entities based on a comprehensive chemical ontology. The classification process involved several stages: (1) preprocessing to ensure data consistency and readiness, (2) feature extraction to identify key structural attributes, (3) rule-based category assignment to classify molecules into appropriate taxonomic categories, and (4) selection of the direct parent category to provide hierarchical context. The output of ClassyFire was generated in SDF format, facilitating subsequent analysis and integration of the classification results into the broader dataset. This method enabled a thorough exploration of the chemical diversity among the genotoxins.

### *In Silico* Biotransformation of Genotoxins

The compiled genotoxin molecules were subjected to *in silico* biotransformation using Biotransformer 3.0 ^21^ to predict phase I (P1), phase II (P2), and enzyme commission (EC)-based metabolism. Biotransformer employs a hybrid approach that integrates rule-based and

machine-learning methodologies to predict the metabolic pathways of small molecules. For each genotoxin, predictions were made for P1, P2, and EC-based pathways independently across three iterations. The ‘cypmode’ parameter was set to “combined,” which incorporates both rule-based and machine-learning techniques. The iterative biotransformation process involves applying the selected *in silico* reactions for a specified number of cycles; for example, products from an initial reaction serve as substrates for subsequent biotransformations in two iterations. SMILES notations of the genotoxins were used as input for the Biotransformer, and the results were generated in SDF format, which facilitated further analysis and integration of the predicted biotransformation products into the dataset.

### Systematic Characterization of Genotoxins

The systematic characterization of genotoxins within chemical space was performed to elucidate their chemical properties. Extended Connectivity Fingerprints (ECFP6) ^40^ were computed for both biotransformed and non-biotransformed genotoxin groups using the RDKit library ^41^. The fingerprints had a dimensionality of 2048 with a radius of 3. Principal Component Analysis (PCA) was then conducted to reduce the dimensionality of the dataset, utilizing the decomposition.PCA function from the Scikit-learn library ^42^ to extract three principal components. The PCA results were visualized using the Matplotlib library, which facilitated the identification of clustering patterns and relationships within the data. This visualization provided insights into the distribution of genotoxins in chemical space, highlighting both biotransformed and non-biotransformed compounds and offering a deeper understanding of their chemical properties and potential interactions.

### Functional Group Analysis

To identify and categorize the predominant functional groups within biotransformed and non-biotransformed genotoxins and to enhance our understanding of their chemical composition, an in-depth functional group analysis was performed using the ChemmineR library ^43^ (v3.46.0). Initially, the genotoxin datasets were converted to the SDFset format, which facilitated the extraction of functional group counts into a matrix. The functional group data were then scaled and visualized using heatmaps to illustrate the distribution and relative abundance of these groups across the datasets. This comprehensive analysis provided detailed insights into the chemical diversity and functional group composition of the genotoxins, offering a deeper understanding of their biochemical properties and interactions.

### Electrophilicity and Oxidative Stress Prediction

Electrophilic potential and oxidative damage are key indicators of genotoxicity. Electrophilic molecules are more likely to attack nucleophilic sites on DNA, leading to DNA adduct formation. To evaluate these properties in biotransformed genotoxic metabolites, we employed the electrophilicity and oxidative stress prediction models within the Metabokiller package ^22^. Metabokiller is an ensemble carcinogenicity prediction tool that quantitatively assesses carcinogenic risk by evaluating various factors, including electrophilicity, cell proliferation potential, oxidative stress, genomic instability, epigenetic alterations, and anti-apoptotic responses. The electrophilicity and oxidative stress models in Metabokiller utilize a multilayer perceptron algorithm. For this analysis, we set the probability threshold for both electrophilicity and oxidative stress at 0.5. Canonical SMILES representations of biotransformed genotoxin products were used as input for Metabokiller, allowing us to systematically predict and categorize their electrophilic and oxidative stress potential. This approach provided valuable insights into the genotoxic properties and potential biological impacts of the metabolites.

### De Novo Generation of DNA Adductome Using MIMOSA

To facilitate the de novo generation of a DNA adductome from genotoxins and their biotransformed products (iterations *k* = 1-3), we employed the Graph Neural Network (GNN) framework of MIMOSA ^23^ (Multi-constraint Molecule Sampling for Molecule Optimization). The initial step in MIMOSA involves generating a vocabulary of molecular substructures, including atoms and rings, from the input molecules. This vocabulary was created by identifying frequent substructures present in both the original genotoxins (iteration *k*=0) and their biotransformed products (iterations *k*=1−3). Substructures appearing in more than ten molecules were included in the vocabulary. For molecular dataset preparation, SMILES notations were loaded from a text file and divided into training (90%) and validation (10%) sets. The Graph Convolutional Network (GCN) model was constructed with 50 input features per node, 100 hidden units, and three graph convolutional layers. The training procedure involved five epochs, with validation checks performed every 5000 iterations. The trained model was saved, and the entire process was executed on a CPU. MIMOSA employs three fundamental molecular operations such as add, replace, and delete, within a Monte Carlo Markov Chain sampling optimization framework. To generate synthetic DNA adducts deoxyribonucleosides (dA, dT, dG, and dC) were used as seed molecules. The generation process involved a population size of 800 across 1200 generations, enabling a thorough exploration of potential DNA adducts derived from both genotoxins and their biotransformed products. Generated DNA adduct structures were evaluated for chemical validity using RDKit ^41^ by converting SMILES representations into molecular objects. Structures that failed SMILES-to-molecule conversion or exhibited valence errors were considered chemically invalid and subsequently removed. The remaining chemically valid structures were retained for further analysis.

### Bioactivity assessment of *in silico*-generated DNA Adducts

To evaluate the bioactivity of *in silico*-generated DNA adducts, we computed bioactivity descriptors using the Signaturizer package ^44^ version 1.1.1. Principal Component Analysis (PCA) was then performed using the decomposition.PCA function from the Scikit-learn library ^42^ to reduce dimensionality. We focused on three principal components to simplify the analysis. The resulting PCA coordinates were visualized with Matplotlib to assess clustering patterns within the bioactivity space of the generated adducts. This analysis provided insights into the bioactivity profiles of the DNA adducts, highlighting distinct clusters that correspond to varying levels of bioactivity.

### Prediction of ESI MS/MS Spectra for DNA Adducts

To predict electrospray ionization (ESI) (+) MS/MS spectra for generated DNA adducts, including precursor [M+H]+ ions, Experimentally Validated Adducts (EVA), and Suspected Adducts (SA), we utilized the CFMID 4.0 Docker image ^26,27^. Predictions were made across collision energies of 10 eV (E0), 20 eV (E1), and 40 eV (E2) using standard parameters provided by the tool. CFMID 4.0 employs a two-step sequential fragment transition model to predict MS/MS spectra, incorporating ring cleavage modeling to improve feature consistency. The tool generates fragmentation graphs where ion fragments are represented as child nodes while neutral fragments are excluded. An adjacency matrix is used to depict molecular topology, and heuristic labeling is applied to fragments. CFMID 4.0 integrates SMIRKS-based fragmentation rules for modular compounds, utilizing a hybrid approach that combines rule-based (MSRB) and machine learning-based (MSML) methodologies. This approach enhances the accuracy and efficiency of spectral predictions, covering a broad spectrum of chemical groups and adduct forms.

### Characterization of Physicochemical and Topological Properties of DNA Adducts

The physicochemical and topological properties of DNA adducts generated by MutAIverse, along with EVA and SA, were systematically characterized using a comprehensive suite of molecular descriptors. Key properties analyzed included atom count, molecular weight, logP (octanol-water partition coefficient), topological polar surface area (TPSA), Kier-Hall index, Kappa indices (Kappa 1, 2, and 3), and Labute’s Approximate Surface Area (ASA). These descriptors were computed using the RDKit library ^41^, providing detailed insights into the molecular features and behaviors of the adducts. This thorough characterization facilitates an in-depth analysis of their chemical properties and potential biological interactions.

### Validation of MutAIverse-Generated DNA Adducts

To validate the DNA adducts generated by MutAIverse, derived exclusively from genotoxins and their biotransformed products, we evaluated their similarity to EVA and SA. This was achieved by comparing the MS/MS spectra of EVA and SA against the MutAIverse spectral library. Cosine similarity scores were calculated using the matchms ^28^ cosine greedy algorithm, with an m/z tolerance set to 0.1 Da. This analysis assessed the congruence between the generated DNA adducts and those experimentally observed, thereby confirming the robustness and accuracy of the MutAIverse framework.

### Spectral Complexity Analysis

A comparative analysis of the spectral complexity of MutAIverse-generated DNA adducts, EVA, and SA was conducted using spectral entropy. Spectral entropy quantifies the information content of a spectrum, reflecting its complexity as described by Shannon entropy in information theory. It is computed based on the number of peaks weighted by their intensity, with typical values ranging from 0 to 5 for small molecule spectra ^45^. Spectral entropy analysis was performed using the ms-entropy package ^46^, version 1.1.3. The MS spectra intensities were normalized such that their sum equaled 1, utilizing the clean spectrum parameter. Entropy similarity was calculated between the MS/MS spectra of EVA and MutAIverse, as well as between SA and MutAIverse, with a default tolerance parameter of 0.05 Da. For each query spectrum from EVA and SA, the maximum entropy similarity was recorded. This approach provided a detailed comparison of spectral properties, validating the fidelity of MutAIverse in accurately representing experimentally observed DNA adduct spectra.

### Spectral Matching with the Mapper Module

The Mapper module is designed to facilitate the spectral matching of MS/MS data from adductomics samples against the MutAIverse spectral library, as well as the EVA and SA libraries. It employs two complementary approaches for assessing spectra similarity: peak matching combined with cosine similarity and Approximate Nearest Neighbour (ANN) search using Hierarchical Navigable Small World (HNSW) graph indexing ^47^. In the first approach, spectral data preprocessing involves normalizing the raw spectra from both the adductomics file and the spectral libraries. Following normalization, the query spectra are compared to the library spectra through peak matching and cosine similarity ^28^ assessments. This method identifies potential matches by evaluating the similarity of spectral peaks. The second approach involves encoding both the query and library spectra into spectral embeddings using a pre-trained word2vec model from FastEI ^48^. These embeddings are indexed in the HNSW graph, which enables efficient vector search. The vector search identifies the nearest neighbor or most similar spectrum to the query spectra within the spectral embeddings. The output of the spectral matching process includes the top nearest neighbor result, along with density plots and histogram visualizations. This integrated framework combines traditional peak matching with cosine similarity and advanced vector-search techniques, ensuring both rapid and accurate identification of spectral matches **(Supplementary** Figure 2e).

### Visualization Using TMAP

TMAP (Tree-based Minimum Spanning Tree Analysis) provides an advanced approach for visualizing high-dimensional datasets by mapping them into a two-dimensional minimum-spanning tree. This method is noted for its superior preservation of both local and global structures compared to traditional dimensionality reduction techniques like t-SNE and UMAP ^25^. TMAP is particularly effective for analyzing extensive chemical datasets, offering clarity and precision in visualization.

In our analysis, MinHash fingerprints (MHFP6, with a dimensionality of 2068) ^49^ were computed from the SMILES representations of MutAIverse DNA adducts using the MHFP encoder package. These fingerprints encapsulate the chemical features of the adducts. The TMAP visualization was then created using the *tmap* package, highlighting various chemical properties such as logP (octanol-water partition coefficient), molecular weight, and ring counts. This approach facilitated a comprehensive and interpretable analysis of the chemical dataset, revealing insights into the distribution and relationships of the DNA adducts.

### AdductLinker Module for Parent Genotoxin Identification

The AdductLinker module is designed to trace the parent genotoxin molecule associated with a detected DNA adduct. The process begins with preprocessing, where the modified segment of the DNA adduct is isolated by extracting the nitrogenous base through maximum common substructure analysis. This step focuses exclusively on the modified portion of the deoxyribonucleoside. After preprocessing, the module generates Extended Connectivity Fingerprints (ECFP6) ^40^ for the extracted modified segment. During the backtracing phase, the k-Nearest Neighbors (k-NN) search algorithm is employed to identify fingerprints in the MutAIverse object that are similar to those of the modified segment. The MutAIverse object contains comprehensive storage of Phase 1 and Phase 2 reaction data for a wide range of genotoxins. The module then determines the Maximum Common Substructure (MCS) ^50^ of the nearest neighbor found in the Phase 1 and Phase 2 reaction data. The MCS is used to retrieve the parent genotoxin molecule from the database. The final output includes detailed information such as the structures of the identified DNA adduct and intermediate metabolite, as well as the parent genotoxin molecule. Additionally, a confidence score is provided, reflecting the probability or degree of confidence in the association between the DNA adduct and its genotoxic parent. This robust backtracing capability offers valuable insights into the biochemical pathways and interactions that lead to DNA adduct formation.

### MutAIverse Python Package

The MutAIverse package offers a comprehensive solution for adductomics research by integrating several key functionalities. It enables the screening of spectral data from adductomics samples against spectral libraries of EVA, SA, and MutAIverse-generated DNA adducts through its Mapper module. Following this, the package facilitates the backtracing of parent genotoxins using its AdductLinker module. This integrated approach streamlines the process of identifying and characterizing DNA adducts and their parent genotoxins, providing a robust toolset for adductomics research.

### External Validation of MutAIverse

External validation of MutAIverse was performed using wSIM-MS/MS (wide selected ion monitoring) data from tumor-adjacent prostate tissue samples of prostate cancer patients (n=4) (RTPS−09−T1: patient 1, RTPS−30−T1: patient 2, RTPS−38−T2: patient 3, and RTPS−41−T2: patient 4) and rat liver tissue samples (control=3, treated=3) exposed to carcinogens/genotoxins, including Benzo[a]pyrene (B[a]P), 4-aminobiphenyl (4-ABP), 2-amino-1-methyl-6-phenylimidazo[4,5-b]pyridine (PhIP), 9H-pyrido[2,3-b]indol-2-amine (AαC), and 3,8-dimethylimidazo[4,5-f]quinoxalin-2-amine (MeIQx), as described in the study by Walmsley et al^29^. The spectral data from these samples were projected against the MutAIverse, EVA, and SA spectral libraries using the Mapper module of the MutAIverse package. Cosine similarity scores were computed to assess the match between experimental spectra and library spectra. The AdductLinker module was then used to trace back the parent genotoxins associated with the detected DNA adducts from the EVA, SA, and MutAIverse-generated adduct libraries. This process validated the MutAIverse framework’s accuracy and robustness in identifying and characterizing DNA adducts.

### Optimization of DNA isolation and hydrolysis from HNC tissue samples

Tissue biopsy samples were collected from head and neck cancer (HNC) patients with a history of smokeless tobacco consumption, including tumor tissue (n=27) and tumor-adjacent control tissue (n=11). Following genomic DNA isolation and enzymatic hydrolysis, the samples underwent untargeted metabolomics analysis. DNA was isolated as given in the protocol of the manufacturer (Nucleospin tissue XS). After the DNA was isolated, purity was checked using a Nanodrop instrument. 1μg human HNC tissue DNA (in 30μl water) was digested. In brief, the DNA sample was denatured at 100°C for 3 min. The sample was cooled rapidly on ice. Later, 3 μL of ammonium acetate (0.1 mol/L, pH 5.3) was added, followed by two units of nuclease P1. The resultant mixture was incubated at 45°C for 2 hours. After, the pH of the mixture was adjusted by adding 4.1µL of freshly prepared ammonium bicarbonate (1 mol/L), and 0.002 units of venom phosphodiesterase were added. The mixture was incubated at 37°C for 2 hours. After incubating with 0.5 unit of alkaline phosphatase at 37°C for 1 hour, the resultant mixture was evaporated and reconstituted with 40µL of methanol: water (1:1) and injected into LC-ESI-QTOF-MS for the analysis of DNA adducts.

### LC-ESI-QTOF-MS analysis of DNA Adducts

The hydrolyzed DNA samples were subjected to liquid chromatography (Agilent 1290 Infinity II HPLC) coupled with high-resolution mass spectrometry (Agilent 6545 LC-ESI-QTOF) in positive ion mode with Agilent dual jet stream electrospray ionization for untargeted determination of DNA adducts in the HNC tissue samples. The mass parameters were set as follows: The source temperature and drying gas flow were maintained at 320°C and 8 L/min, respectively. The nebulizer pressure was set to 35psi. The sheath gas temperature and sheath gas flow rate were set to 350 °C and 11 L/min, respectively. The capillary voltage was set to 3500 V, and the nozzle voltage was set to 1000V. The data was acquired in MS mode with a mass range of 50-1700 m/z. The MS acquisition rate and time were set to 1 spectra/s and 1000 ms/spectrum, respectively. The MS data was acquired using MassHunter workstation software. The DNA adducts were separated using Agilent Zorbax RRHDC Eclipse plus C18 column (2.1 ×100 mm, 1.8µm) maintained at 40°C using mobile phases A: 0.1% formic acid in milliQ water and mobile phase B: 0.1% formic acid in acetonitrile. The injection volume and flow rate were set to 10µL and 0.3mL/min, respectively. The following gradient elution mode was used for the separation of the compound: 0 min: 90%A, 3 min: 85%A, 5 min:78%A, 10 min: 60%A, 13 min: 50%A, 15 min: 30%A, 18-20 min: 90%A with total run time of 20 minutes.

### DNA Adductomics on tissues from head and neck cancer patients

Raw data files (.d files) were converted to mzML format and underwent peak centroiding using the msconvert utility of ProteoWizard ^51^. These converted files were then searched against the MutAIverse adductome library, as well as the EVA and SA DNA adduct libraries, to compute cosine similarity scores. The AdductLinker module was employed to trace back the parent genotoxins associated with the detected EVA, SA, and MutAIverse-generated DNA adducts. Top hits, defined as those with a cosine similarity greater than 0.9, were identified, and further, their precursor m/z values were confirmed by high-resolution mass spectrometry.

### Multivariate analysis of samples

To assess variation across samples based on their spectral composition, we conducted a multivariate analysis using MetaboAnalyst 6.0 ^52^. This platform employs peak detection, peak picking, peak alignment, and gap-filling algorithms to identify and standardize features across samples. Principal Component Analysis (PCA) was then performed to explore clustering patterns and visualize the relationships among the samples.

### DNA Adduct Abundance and Patient Age Analysis

Intensity values for m/z 377.18 and 266.07 in tumor and tumor-adjacent control samples were extracted from the feature list generated via LC-MS processing using MetaboAnalyst 6.0 ^52^. To assess statistical significance between different sample types, the Mann-Whitney U test was performed. Additionally, to analyze the abundance of m/z 377.18 and 266.07 relative to patient age, Pearson’s correlation coefficient was calculated along with p-values for linear regression analysis.

### Statistical Analysis

All statistical analyses were performed using Scipy Python package (version 1.14.0). Kolmogorov-Smirnov test was used to test differences in probability distributions for statistical significance. Mann-Whitney U test was performed to compare the medians of two non-parametric distributions. The p-value cutoff for significance was 0.05. Significance levels are: * < 0.05, ** < 0.01, *** < 0.001, and **** < 0.0001.

### Code Availability

The source code for MutAIverse is available on the GitHub repository (https://github.com/the-ahuja-lab/MutAIverse). Python package for MutAiverse is available via pip: https://test.pypi.org/project/MutAIverse. We also set up the entire run code on Google Colab: https://colab.research.google.com/drive/1cidZQgZHD0wDcTsRMbO2jY0NFZfKW-Ig?usp=sharing.

### Ethical Clearance

The local Medical Ethics Committee at Dr. Bhubaneswar Borooah Cancer Institute, Department of Atomic Energy, Government of India (Reference number: BBCI-TMC/Misc-01/MEC/252/2021) approved the use of head and neck tumor tissues from patients. This approval was in accordance with the guidelines of the Indian Council of Medical Research, New Delhi, Government of India.

### Declaration of Interests

The authors declare no competing interests.

## Acknowledgments

The authors thank the IT-HelpDesk team of IIIT-Delhi for assisting with the computational resources. We thank all the members of the Ahuja lab for their intellectual contributions at various stages of this project. The Ahuja lab is supported by the Ramalingaswami Re-entry Fellowship (BT/HRD/35/02/2006), a re-entry scheme of the Department of Biotechnology, Ministry of Science & Technology, Government of India, and an intramural Start-up grant from Indraprastha Institute of Information Technology-Delhi. Borkar lab is funded by the Science and Engineering Research Board (SERB), Ministry of Science and Technology, Government of India, New Delhi, under the project titled “Identify the DNA Adduct and Associated Metabolic Alterations in Upper Aerodigestive Tract Cancer with Smokeless Tobacco Chewers in the Northeast Region of India: A Metabolomics Approach (EEQ/2020/000393)”.

## Author Contributions

The study idea was conceived by G.A. Computational and experimental parts of the manuscript were designed and supervised by G.A. and R.M.B., respectively. DNA adductomics was performed by S.J. under the supervision of R.M.B. MutAIverse workflows were designed by G.A. S.S., and S.K.M designed, and the architecture was built by S.S., S.K.M, and A.V. Data analyses were performed by S.S., S.K.M., S.J., A.V., Sa.S., V.G., S.A., A.M., A.S., S.C., S.D., S.K., J.T., A.M., D.S, and G.A. Human samples collections were performed by A.R., A.D., K.K., K.D., and A.S, and processed by S.J., and R.M.B. Illustrations were drafted by S.S., and S.K.M. G.A. and R.M.B wrote the paper. All authors have read and approved the manuscript.

## Supplementary Tables

**SI Table S1:** Tabular representation containing detailed information about the sources of genotoxins.

**SI Table S2:** Tabular representation containing detailed information about the chemical classification of genotoxins.

**SI Table S3:** Tabular representation containing detailed information about the substructure frequency of biotransformed genotoxins across k-Biotransformations (k=0,1,2,3).

**SI Table S4:** Tabular representation containing detailed information about EVA and SA adducts.

**SI Table S5:** Tabular representation containing detailed information of maximum cosine scores for MutAIverse DNA adducts when MS/MS spectra of EVA and SA searched against the MutAIverse spectral library.

**SI Table S6:** Tabular representation containing detailed information about the metadata of head and neck cancer patients.

## Notes

### Competing Interest Statement

The authors have declared no competing interest.

