## Supplementary figures and images for "MutAIverse: An AI-Powered, Mechanism-backed Platform for Discovering Novel DNA Adducts and their precursor Genotoxins"

### Supplementary Figure 1

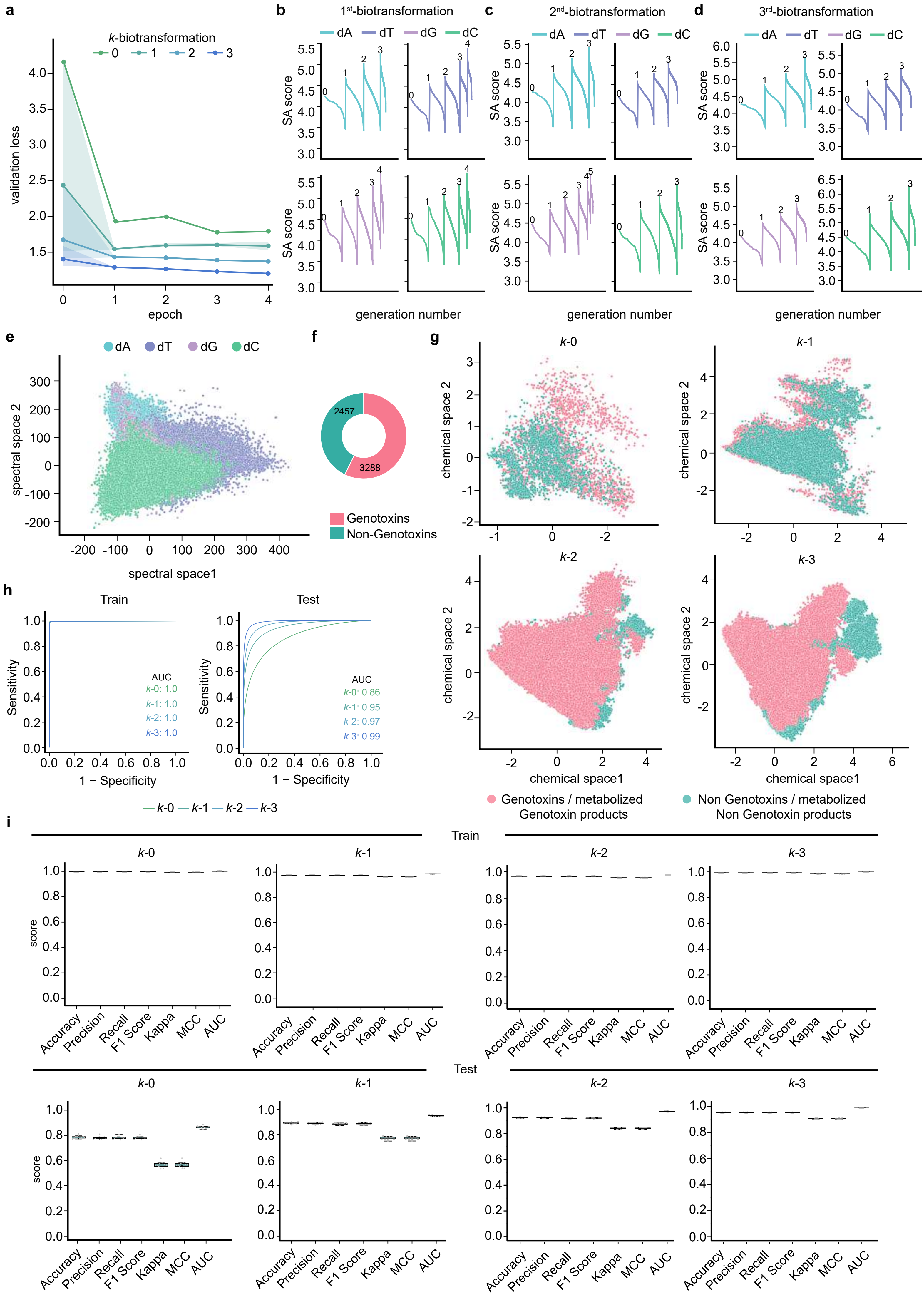


Supplementary Figure 1

### Supplementary Figure 2

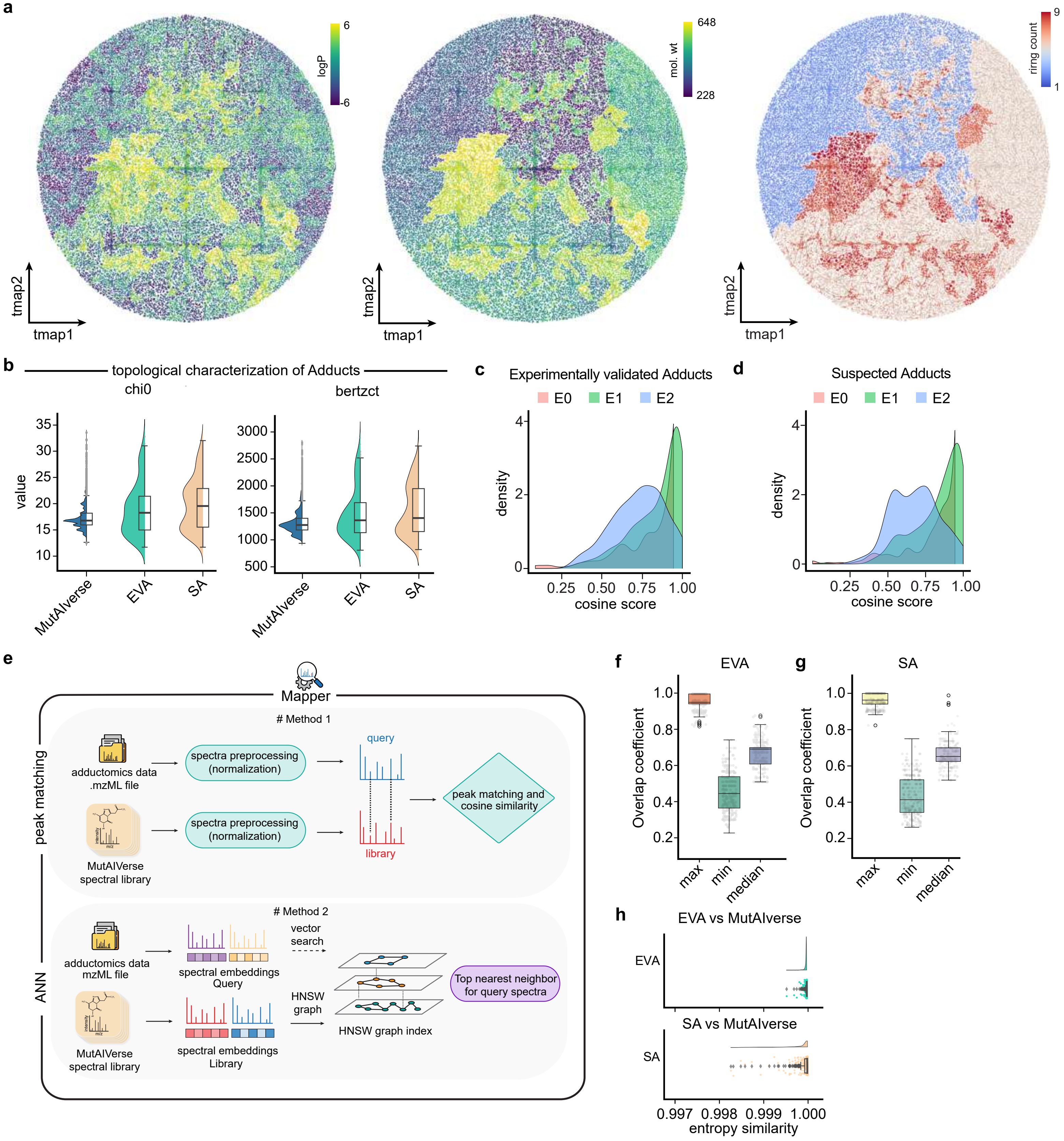


Supplementary Figure 2

### Supplementary Figure 3

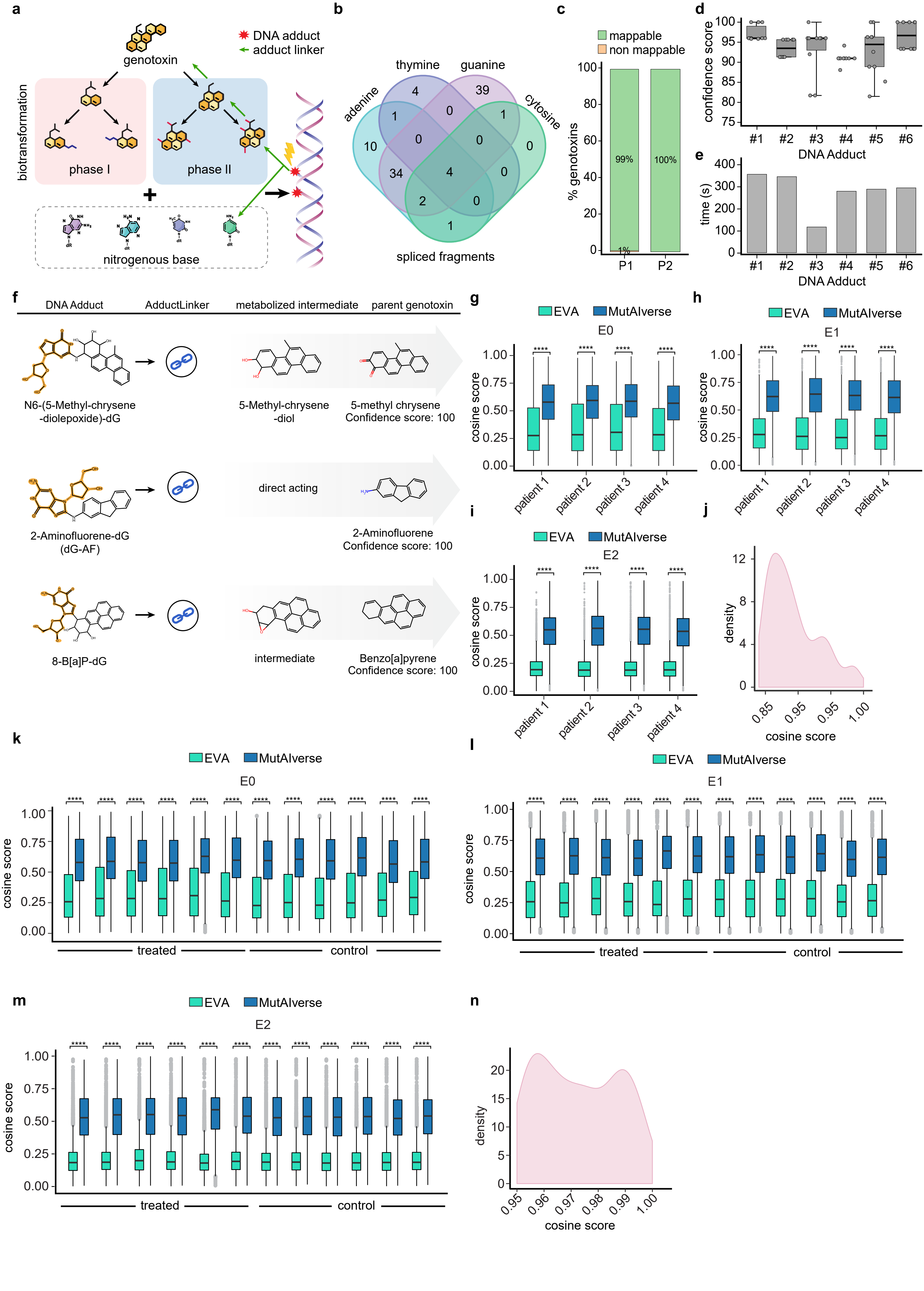

Supplementary Figure 3

### Supplementary Figure 4

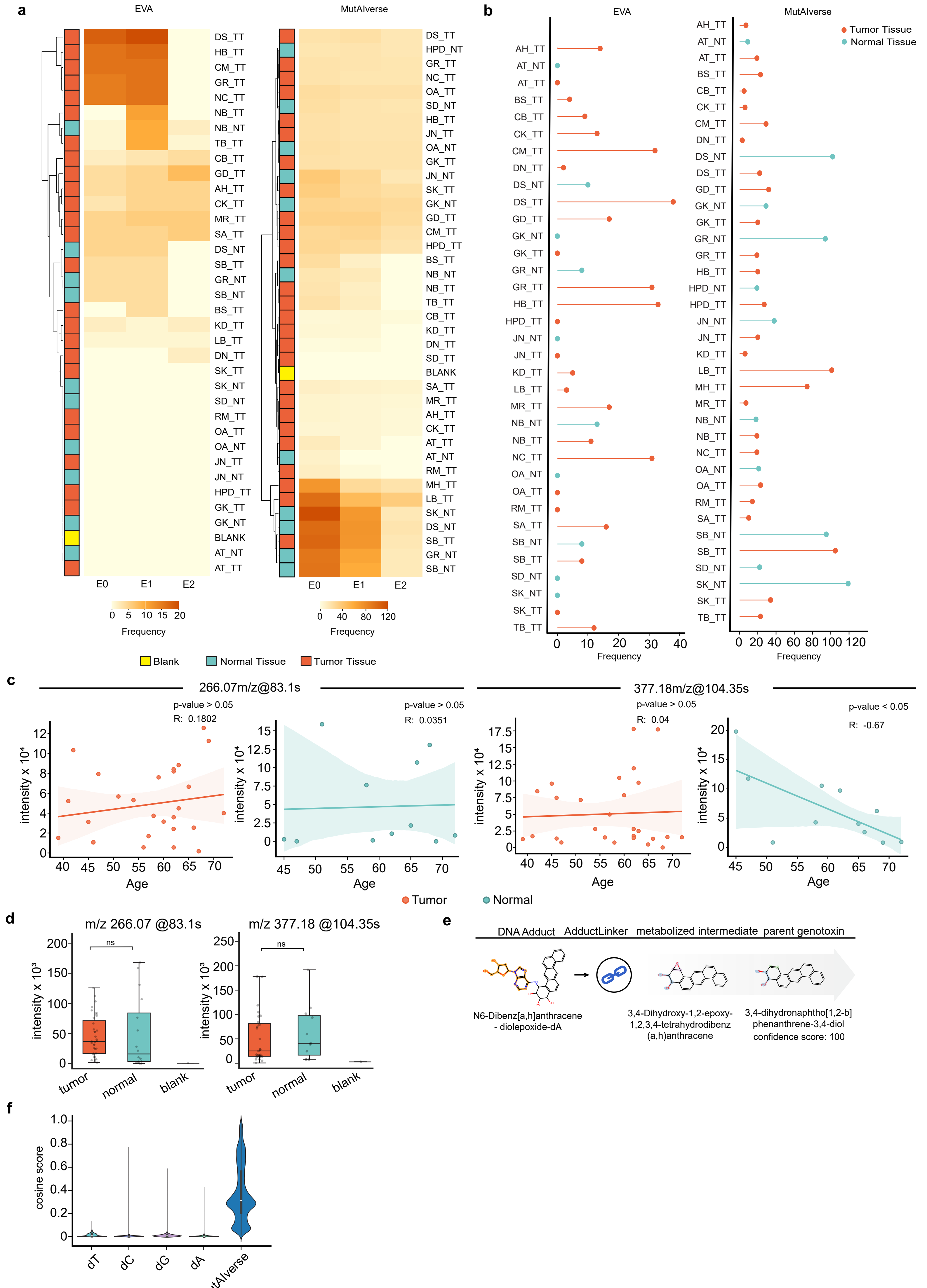


Supplementary Figure 4
